# High density cultivation for efficient sesquiterpenoid biosynthesis in *Synechocystis* sp. PCC 6803

**DOI:** 10.1101/834499

**Authors:** Dennis Dienst, Julian Wichmann, Oliver Mantovani, João Rodrigues, Pia Lindberg

**Author notes:** Correspondence should be addressed to Pia Lindberg.

## Abstract

Cyanobacteria and microalgae are attractive phototrophic host systems for climate-friendly production of fuels and other high-value chemicals. The biosynthesis of an increasing diversity of industrially relevant compounds such as terpenoids has been demonstrated in recent years. To develop economically feasible and sustainable process designs, major challenges still remain regarding intracellular carbon partitioning, specific metabolic pathway activities and efficient cultivation strategies. Here, we present a technical study on comparative characteristics of sesquiterpene and sesquiterpene alcohol accumulation in engineered strains of *Synechocystis* sp. PCC 6803 (substrain GT-U) under different growth conditions and cell densities. This study particularly focuses on the basic applicability of a commercial High Density Cultivation platform in the presence of a dodecane overlay, which serves as a standard *in-situ* extractant and sink for various hydrophobic biochemicals. Significantly, the presented data demonstrate high volumetric productivities of (*E*)-α-bisabolene under high-density conditions that are more than two orders of magnitude higher than previously reported for cyanobacteria. Operating in a two-step semi-batch mode over a period of eight days, average final volumetric titers of 179.4 ± 20.7 mg * L^−1^ were detected. Likewise, the sesquiterpene alcohols (-)-patchoulol and (-)-α-bisabolol accumulated to many times higher levels in high density cultivation than under standard batch conditions, with final titers of 17.3 ± 1.85 mg * L^−1^ and 96.3 ± 2.2 mg * L^−1^, respectively. In contrast, specific product accumulation (mg * L^−1^ * OD_750_^−1^) was compromised particularly for bisabolene in the high density system during phases of high biomass accumulation rates. Volumetric productivities were high during linear growth at high densities, distinctly outperforming standard batch systems. While the presented data highlight the benefits of high-density strategies for highly efficient phototrophic terpenoid production, they further point at the presence of major metabolic bottlenecks for engineered terpenoid biosynthesis and the requirement for systematic and/or targeted strategies to sustainably redirect inherent carbon fluxes in cyanobacteria. Together, our data provide additional insights into growth- and density-related effects on the efficiency of product accumulation, introducing low-scale High Density Cultivation as a rapid and efficient platform for screening of heterologous terpenoid production in cyanobacteria.

## Introduction

Cyanobacteria are oxygenic phototrophs with a versatile metabolism and therefore constitute an economically appealing platform for the sustainable production of a diversity of industrially relevant chemicals. In particular, the natural occurrence of the methylerythritol-4-phosphate (MEP) pathway [1] allows the engineered biosynthesis of a diversity of high-value terpenoid compounds such as pharmaceuticals, cosmetics, and biofuels [reviewed in 2, 3]. While numerous pioneering proof-of-concept studies already demonstrated the heterologous biosynthesis of isoprene (C_5_), mono-(C_10_), sesqui-(C_15_), di-(C_20_) and tri-(C_30_) terpenes in cyanobacterial host strains [4–9], the reported titers of these compounds are typically two to three orders of magnitudes lower than those of e.g. alcoholic fuels [cf. overviews in 3, 10]. The major limitation of these processes arise from the inherent carbon partitioning characteristics, which favor primary metabolism and biomass accumulation, resulting in weak carbon fluxes through the native MEP pathway [11]. In accordance, a strategy to bypass these inherent flux limitations by heterologous expression of the complete bacterial mevalonate (MVA) pathway in *Synechocystis* sp. PCC 6803 (from here: *Synechocystis*) resulted in 2.5-fold higher isoprene yields [12]. Likewise, numerous rational design studies targeting selected bottleneck steps within the MEP pathway as well as selected steps of upstream carbon fluxes lent support to this concept [8, 9, 13–19]. Further strategies sought to challenge the problem of inherently weak terpene synthase activities by increasing cellular enzyme titers, concurrently pushing carbon flux towards the heterologous metabolic sink. To this end, major improvements were reported by systematic, genetic optimization of terpene synthase expression cassettes, and/or by designing highly expressed fusion proteins [13–15, 20, 21].

Aside from these specific metabolic and physiological bottlenecks, general drawbacks of photoautotrophic production strategies arise from inherent technical limitations of the established cultivation systems, particularly regarding the supply of essential nutrients and light [22, 23]. These limitations include inefficiencies in mass transfer of gaseous CO_2_ to liquid growth medium, as well as physiological constraints due to rapid and distinct alkalization of the growth medium, particularly when using bicarbonate salts as inorganic carbon source [24].

Classical screening setups for terpenoid production in cyanobacteria involve photoautotrophic batch cultivation in shaking flasks or aerated vessels supplied with bubble columns of CO_2_-enriched air. When calculated to standardized time frames of four days, these protocols result in rather dilute cultivation setups as well as low volumetric product titers of up to ~8 mg * L^−1^ or specific titers of up to ~20 mg * g^−1^ DCW, respectively [summarized in 3, 25]. An appreciable exception is a study on production of the volatile compound isoprene in *Synechococcus elongatus* PCC 7942, for which volumetric titers of more than 400 mg * L^−1^ were reported in 4 days [13]. In that particular long-term experiment, the strains were grown under optimized conditions in photobioreactors at 37 °C with a combination of CO_2_ and additional bicarbonate supply at 100 µmol photons * m^−2^ * s^−1^, including daily media replenishment. When grown under standard batch conditions (30 °C, shaking flasks, additional bicarbonate supply, 55 µmol photons * m^−2^ * s^−1^), yields from the same strains were about one order of magnitude lower, pointing at the requirement for a stable and well buffered system for sufficient inorganic carbon supply. Furthermore, individual inherent properties like hydrophobicity, oxygenation and molecular size of terpenoid products can impose specific challenges on common harvesting procedures like organic *in situ* extraction [26].

High density cultivation (HDC) is a recently developed cultivation system that tackles these problems by implementing a membrane-mediated CO_2_ supply technology, combined with optimized nutrient supply and high light intensities [27]. This system facilitates rapid, sustainable biomass accumulation and was demonstrated to further raise the potential for the recovery of natural products from cyanobacteria [28–30].

To assess the potential of HDC for improved terpenoid production yields from cyanobacteria, three industrially attractive sesquiterpenes with different physicochemical properties were selected as reporter compounds: (1) the monocyclic (*E*)-α-bisabolene (C_15_H_26_O, from here: bisabolene), a non-oxidized sesquiterpene found as an intermediate for conifer resin biosynthesis that serves as a precursor of the D2 Diesel-type fuel bisabolane [31]; (2) its oxidized form (-)-α-bisabolol (C_15_H_26_O, from here: bisabolol), a sesquiterpene alcohol derived from chamomile plants with pharmaceutical and cosmetic applications [32]; and (3) the tricyclic sesquiterpene alcohol (-)-patchoulol (C_15_H_26_O, from here: patchoulol), which is the major odorant in patchouli oil and precursor for synthesis of the chemotherapeutic agent taxol [33, 34]. These compounds can be directly generated by heterologous expression of each a single terpene synthase (TS) gene, the products of which catalyze the cyclization of the common C15 precursor farnesyl pyrophosphate (FPP).

Photoautotrophic production of bisabolene was previously demonstrated in engineered strains of *Synechococcus* sp. PCC 7002 [35] and the green alga *Chlamydomonas reinhardtii* [36], yielding final titers of 0.6 mg * L^−1^ (in 4 days) and 3.9 mg * L^−1^ (in 7 days), respectively. Patchoulol titers of 1.03 mg * L^−1^ (in 7 days) were reported for *C. reinhardtii* [37], while heterologous bisabolol synthesis has so far been demonstrated exclusively in heterotroph microbial hosts, *Saccharomyces cerevisiae* and *Escherichia coli* [38, 39]. Here we expand the range of microbial host systems for the production of these compounds to the model cyanobacterium *Synechocystis*, presenting a comparative study on the potential of HDC for photoautotrophic sesquiterpenoid production. We highlight growth phase-dependent product accumulation in different cultivation systems of non-hydroxylated (bisabolene) and hydroxylated (bisabolol, patchoulol) terpenoid products.

## Results and Discussion

### Design and experimental strategy for heterologous biosynthesis of sesquiterpenes

For the implementation of heterologous biosynthesis of bisabolene, bisabolol and patchoulol in *Synechocystis*, we took advantage of the intrinsic FPP precursor pool provided by the native MEP pathway, without any further engineering of upstream pathway sections (Fig. 1A). For ectopic expression of the terpene synthases, codon optimized coding sequences (CDS) of bisabolene synthase from *Abies grandis* (*Ag*BIS, GenBank accession no.: MG052654), (-)-α-bisabolol synthase from *Matricaria recutita* (*Mr*BBS, GenBank accession no.: KJ020282) and patchoulol synthase from *Pogostemon cablin* (*Pc*Ps, GenBank accession no.: KX097887) were each linked to the promoter:RBS module P_*petE*_:BCD2 (from here: P_*petE*_), which mediates strong Cu^2+^-inducible expression (Fig. 1B). The expression module was further linked to a codon optimized version of the fluorescence reporter mVenus [40], which was deployed as isogenic negative control for sesquiterpene accumulation and quantitative expression proxy during all experiments. All expression cassettes were located on the RSF1010-derived, autonomously replicating plasmid pSHDY*in [41, Fig. S1]. After conjugational plasmid transfer and positive clone selection (Fig. 1C), S*ynechocystis* strains were cultivated in liquid medium, according to the protocols for the three different cultivation strategies: (i) 6-well-plate cultures for initial screening; (ii) Multicultivator (MC) cultivation and (iii) HDC with nutrient-enriched media. Production of each terpenoid compound was assessed by GC-MS and GC-FID analysis.

**Figure 1:**
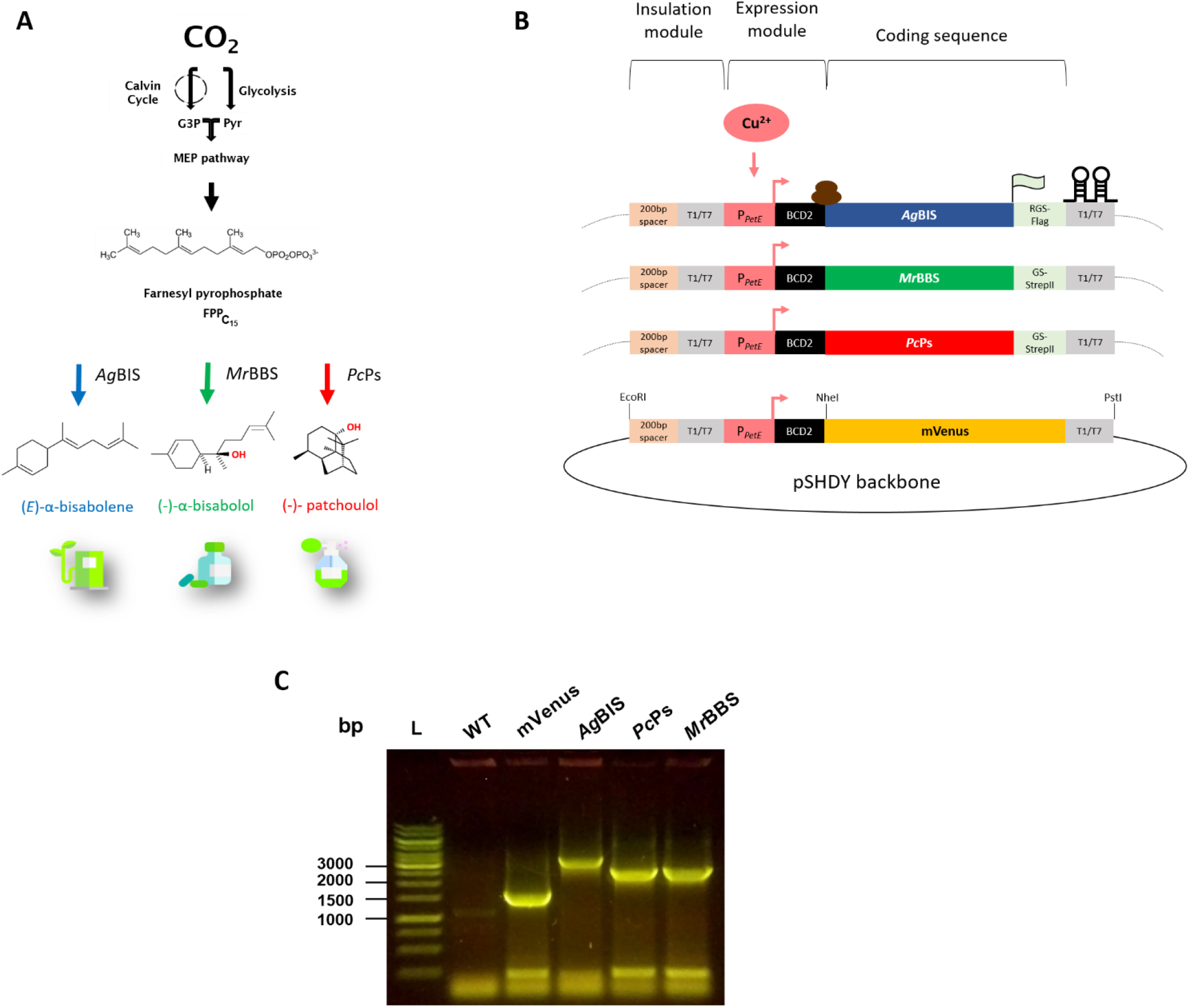
Sesquiterpene pathway design in *Synechocystis*. (A) Pathway for targeted biosynthesis of bisabolene, bisabolol and patchoulol from CO_2_ in *Synechocystis* by introducing genes encoding bisabolene synthase from *A. grandis* (*Ag*BIS, blue arrow), (-)-α-bisabolol synthase from *M. recutita* (*Mr*BBS, green arrow) and patchoulol synthase from *P. cablin* (*Pc*Ps, red arrow), respectively. The common sesquiterpene precursor FPP derives from the native MEP pathway. (B) Schematic representation of expression constructs used for sesquiterpene biosynthesis in the pSHDY*in vector (a derivative of pSHDY [41]). Gene expression is mediated by synthetic expression modules, consisting of the native copper-inducible promoter P_*petE*_[42, 43] linked to the *bicistronic design* BCD2 for reliable translation initiation [44]. The CDS of *Ag*BIS is C-terminally fused to a 1x Flag tag via an RSGSGS linker (RGS); *Mr*BBS and *Pc*Pc are fused to a Strep tag II via a GSGSGS linker (GS). The mVenus is devoid of an epitope tag. All expression cassettes are terminated by the T1/T7 double terminator (BBa_B0015). The dimensions of the genetic parts were chosen for schematic representation of the modular arrangements and do not reflect actual size relations. (C) Confirmation of positives clones of *Synechocystis* harboring plasmids pSHDY*in_PpetE:BCD2:mVenus, pSHDY*in_PpetE:BCD2:AgBIS:Flag, pSHDY*in_PpetE:BCD2:PcPs:Strep and pSHDY*in_PpetE:BCD2:MrBBS:Strep, respectively. Colony PCR reactions were performed with the universal primer pair CC16/ CC103 (cf. Table S1). Fragment sizes are 1615 bp (mVenus), 3374 bp (*Ag*BIS), 2603 bp (*Pc*Ps) and 2654 bp (*Mr*BBS), respectively. FPP, farnesyl pyrophosphate; G3P, glyceraldehyde 3-phosphate; Pyr, pyruvate; WT, wild type. Icons made by Freepik from www.flaticon.com

### Assessment of sesquiterpene biosynthesis in 6 well plates

For the initial screening procedure and expression module assessment, positive clones of each of the four constructs were grown over four days in 6 well plate cultures (Fig. 2A) under constant agitation and moderate illumination. All strains were cultivated both in Cu^2+^-free medium (-Cu) and in medium containing 2µM Cu^2+^ (+Cu), with a dodecane overlay to trap the product.

**Figure 2:**
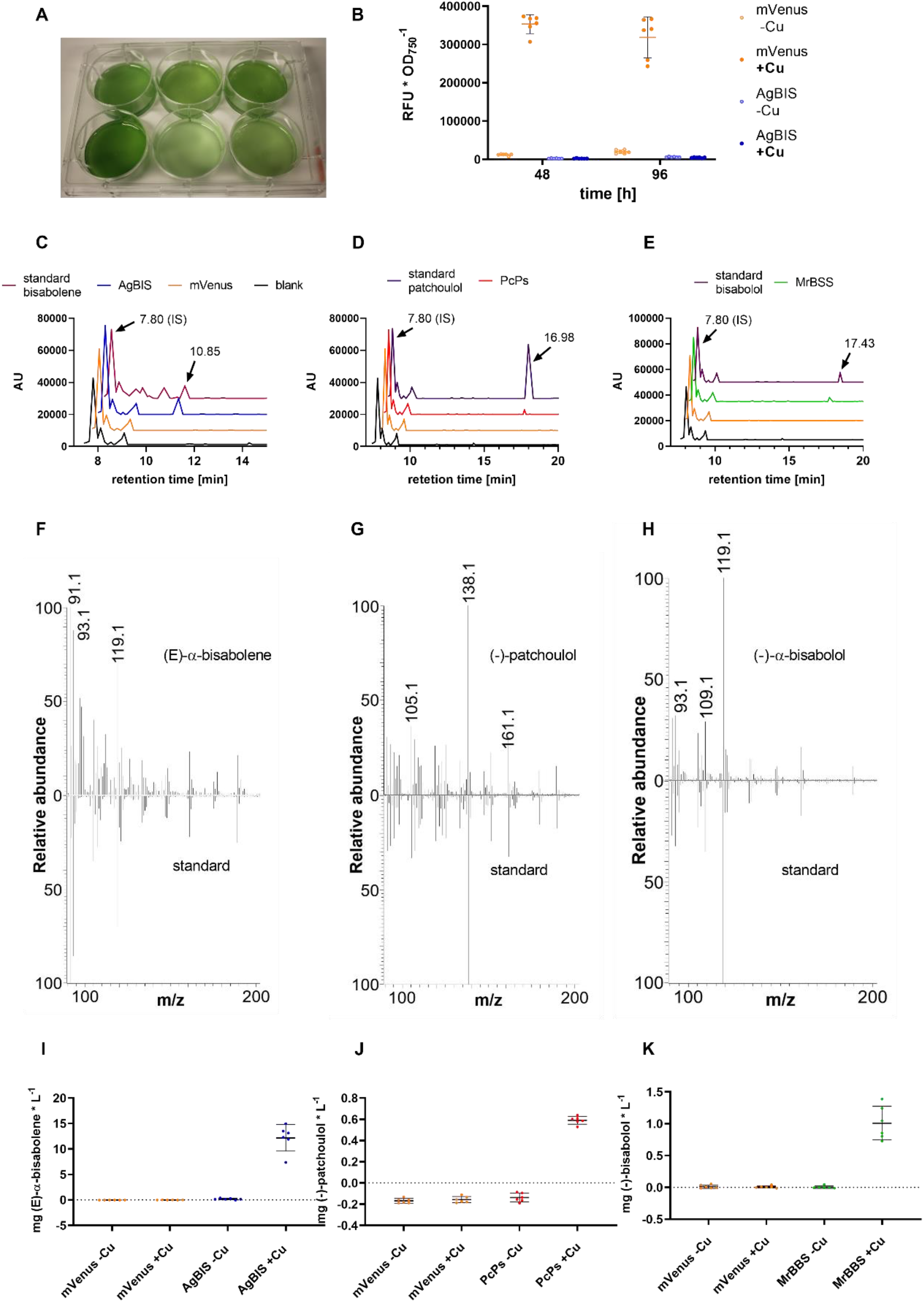
GC-FID and GC-MS detection and quantification of sesquiterpenoids in *Synechocystis*. (A) Strains expressing mVenus, *Ag*BIS, *Mr*BBS and *Pc*Ps were cultivated for 4 days in 6-well plates with 20 % [v/v] dodecane overlay, which was directly subjected to quantitative GC-FID analysis, deploying β-caryophyllene as internal standard (IS). (B) mVenus accumulation profile at time points 48h and 96h from cultures treated with 2 µM CuSO_4_ (+Cu) and non-induced cultures (-Cu). Data from strain *Ag*BIS is shown as negative fluorescence control. Fluorescence data was normalized to the corresponding OD_750_ data and is depicted as mean values of sextuplicates with SD. (C), (D), and (E) GC elution profiles of bisabolene, patchoulol and bisabolol, respectively. Sesquiterpene identity was verified by alignment of retention times to commercial references; overlays from cultures expressing mVenus as well as ‘blank’ dodecane samples (IS only) served as negative controls. Signals at 7.80 min correspond to the IS. Signals at 10.85 min, 17.43 min and 16.98 min correspond to bisabolene, bisabolol and patchoulol, respectively. (F), (G), (H) Mass spectra of bisabolene, patchoulol and bisabolol detected by GC-MS analysis of dodecane samples. Signal spectra obtained with the corresponding reference compound are depicted in the lower panel. (I), (J) and (K) Volumetric titers from induced (+Cu) and non-induced (-Cu) cultures. Quantitative diagrams show individual values from biological sextuplicates and their mean with SD.

Elution profiles in the GC-FID analysis compared with the corresponding commercial standard compounds confirm the presence of bisabolene, bisabolol and patchoulol in the respective strains, while these compounds were absent from samples of control strain P_*petE*__mVenus (Fig. 2C-E). This data is further supported by the mass fractionation patterns derived from the GC-MS analysis, which match with the corresponding reference standard of each compound (Fig 2 F-H). The presence of further isomers in the bisabolene standard (Fig. 2C) was previously reported [36] and correspondingly included in the calculations. Reporter fluorescence readout indicated 30-fold and 16-fold higher expression levels from induced P_*petE*__mVenus constructs compared to the non-induced controls at time points 48h and 96h, respectively (Fig. 2B). Besides the lower apparent increase mediated by the induction, the values measured at time point 96h further show a higher variance, which might result from more variable Cu^2+^ supply at higher cell densities, since no further addition of Cu^2+^ was made. As for the levels of bisabolene, an apparent 70-fold induction was detected at time point 96h (final titer 12.2 ± 2.5 mg * L^−1^), while patchoulol and bisabolol levels were below the detection limit in the non-induced cultures (Fig. 2 I-K). Final titers in the induced cultures were 0.59 ± 0.03 mg * L^−1^ and 1.0 ± 0.24 mg * L^−1^ for patchoulol and bisabolol, respectively. Hence, recovered volumetric product yields for bisabolene appear one order of magnitude higher than those for both hydroxylated sesquiterpenoids under these conditions. Together, these data suggest good correlation between the P_*petE*_:BCD2 expression module activity and product formation. However, slight amounts of both mVenus and bisabolene product were detected also in non-induced cultures. Since all clones were consequently propagated in disposable plastic vessels, basal expression from P_*petE*_:BCD2 is likely due to traces of Cu^2+^ in the water source. In conclusion, under these conditions the P_*petE*_ promoter enables sensitive and strong induction of gene expression, however, exhibits a rather poor dynamic range, in accordance with previous observations [45].

### Sesquiterpenoid production in multicultivator (MC) tubes with bicarbonate supply

To assess kinetic characteristics of time- and growth-dependent sesquiterpene accumulation in a standard batch mode, strains P_*petE*__*Ag*BIS, P_*petE*__*Pc*Ps, P_*petE*__*Mr*BBS and P_*petE*__mVenus were grown in continuously aerated MC tubes (Fig. 3A) with additional bicarbonate supply. Measurement of OD_750_ and mVenus fluorescence were performed daily; dodecane was sampled bidaily. The growth curves depicted in Fig. 3B display typical tripartite profiles as described previously [46]. In this specific setup, growth is preceded by an initial 24 h lag phase (phase 0), which probably results from adaptation to changing media composition and light regime as well as to shear stress during the initial centrifugation of the preparatory cultures [47, 48]. The cultures exhibit exponential growth over a period of four days (phase 1: 24h-120h), followed by a two-day linear growth phase (phase 2: 120h-144h) and eventually a stationary phase (phase 3: 146h – 192 h). After the end of phase 2 (tp: 144 h) cultures were diluted with fresh medium (including 50 mM NaHCO_3_), and light intensities were elevated from 50 to 70 µmol photons * m^−2^ * s^−1^ before cultivation continued from tp 146h. Interestingly, the decrease of growth rates in phase 3 was less pronounced for P_*petE*__mVenus and P_*petE*__*Mr*BBS (Fig. 3C). Constant and stable relative accumulation of mVenus in P_*petE*__mVenus (Fig. 3D) indicates robust activity of the P_*petE*_:BCD2 expression module over the entire cultivation period, confirming that one initial addition of Cu^2+^ is sufficient for sustainably high expression levels under these conditions.

**Figure 3:**
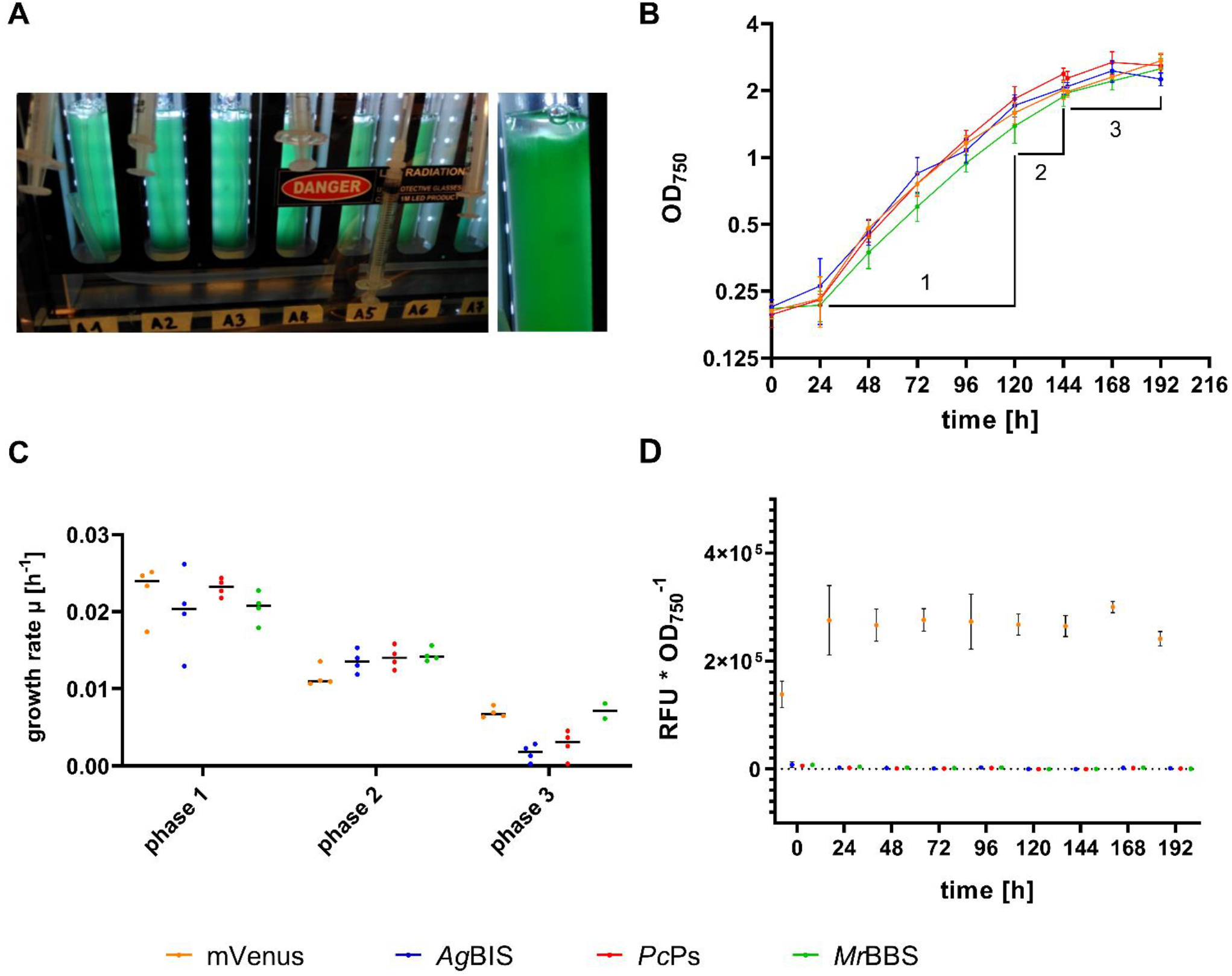
Growth characteristics and heterologous gene expression in MC batch cultures. (A) Batch cultivation setup in MC-1000 reactors (Photon Systems Instruments). Bicarbonate-enriched liquid cultures were supplemented with each 5% [v/v] dodecane overlay and sparged with a constant stream of air. (B) Cell densities (OD_750_) of strains *P*_petE__AgBIS, *P*_petE__*MrBBS*, *P*_petE__PcPc *and P*_petE__mVenus were recorded daily over a time period of 9 days. Data represent mean values from four independent cultures with SD. The y-axis scale is logarithmic (log2). Growth phases are indicated (C) Growth rate µ calculated from the OD_750_ data in (B). Data represent mean (lines) and single (dots) values with SD. Phase 1, 2 and 3 refer to time periods 24-120h, 120-144h and 146-192h, respectively. (D) mVenus accumulation profile over time. Fluorescence data were divided by the corresponding OD_750_ data [λ Ex./Em. 485/535nm * OD_750_^−1^] and are depicted as mean values with SD.

The kinetics of product accumulation showed similar patterns for bisabolol and patchoulol (Fig. 4B, C), while bisabolene accumulation initially appeared delayed in time, but reached higher volumetric and specific titers of 7.4 mg * L^−1^ (Fig. 4A) and 3.3 mg * L^−1*^ OD_750_^−1^ (Fig. 4G) at tp 192h, respectively. By comparison, final patchoulol and bisabolol titers were 1.3 mg * L^−1^ (Fig. 4B) / 0.5 mg * L^−1*^ OD_750_^−1^ (Fig. 4H) and 2.9 mg * L^−1^ (Fig. 4C) / 1.2 mg * L^−1^* OD_750_^−1^ (Fig. 4I), respectively. Notably, in all strains volumetric product titers increase over all three growth phases. In accordance with biomass accumulation, volumetric productivity (mg * L^−1^ *d^−1^) slightly increases over time and the growth phases in all strains (Fig. 4D-F). Specific product yields showed a clear increment for bisabolene and patchoulol (Fig. 4 G-H), while per-cell yields of bisabolol stalled upon entry into phase 3 (Fig. 4 I). The latter effect is presumably resulting from a higher specific growth rate of strain P*petE*:*Mr*BBS in phase 3 compared to P*petE*_AgBIS and P*petE*_PcPs, rendering the cellular carbon partitioning characteristics in a state that favors biomass production over terpenoid biosynthesis.

**Figure 4:**
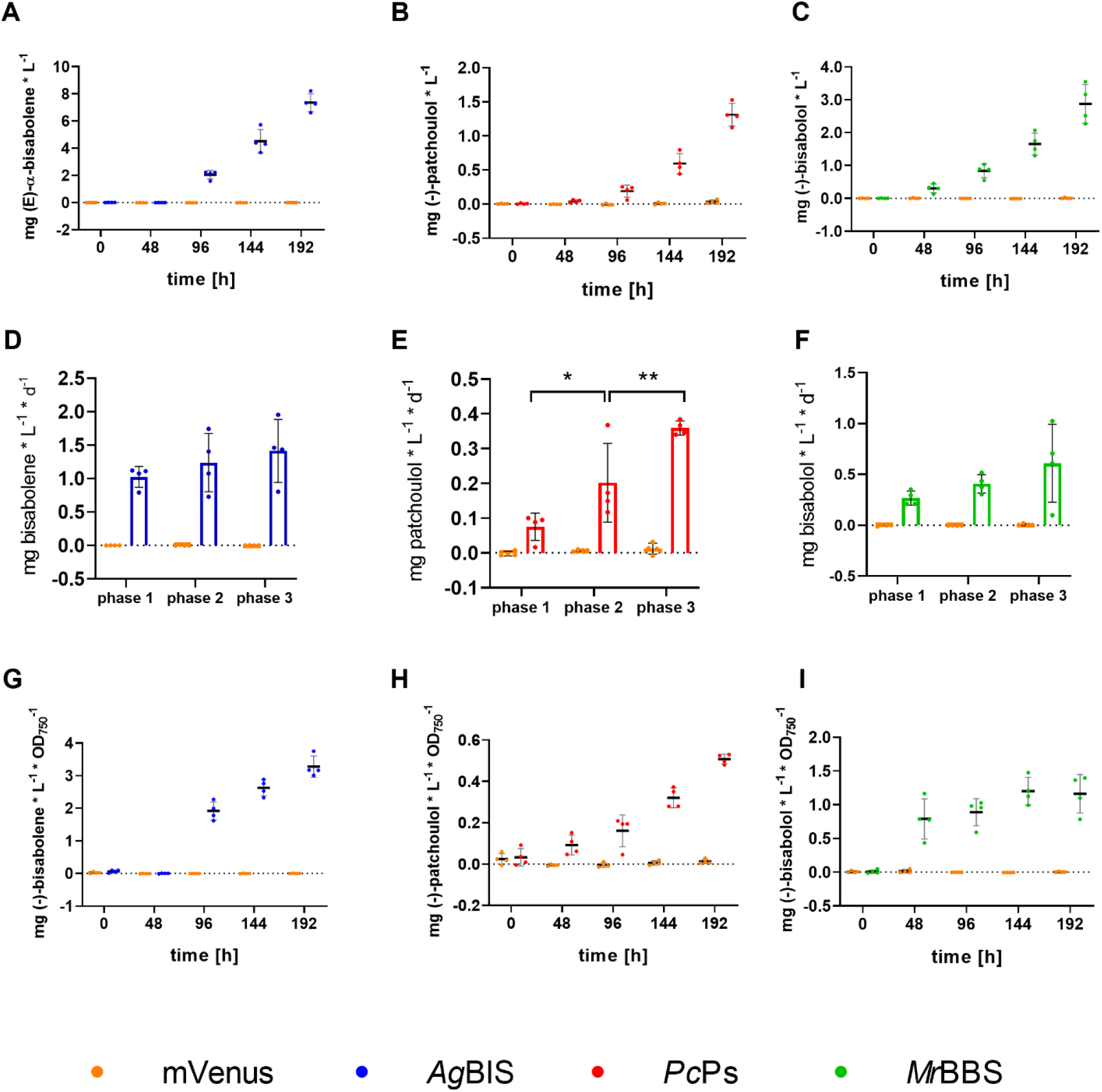
Time-dependent sesquiterpene accumulation profiles from batch cultures grown in MC reactors. (A-C) Volumetric titers of bisabolene, patchoulol and bisabolol, respectively. Quantities were measured by GC-FID directly from the dodecane overlay and normalized to the total culture volume (mg * L^−1^). (D-F) Corresponding volumetric productivities calculated as the temporal yields (mg * L^−1^ * d^−1^) for growth phases 1 (24-120h), 2 (120-144h) and 3 (146-192h), respectively. Statistical analysis was performed using the two-way ANOVA tool of the GraphPad Prism software (version 8.2.1.). *P < 0.05; **P < 0.01. (G-I) Specific product titers calculated as the quotient between volumetric titers and cell densities of the cultures (mg * L^−1^ * OD_750_^−1^).

Together these data demonstrate a trend towards growth phase and/or biomass related dependency of cell culture productivity, suggesting higher cell densities to be beneficial for the process of sesquiterpene production. However, it is of note that the apparent growth-related productivity effects in this data are only statistically significant for patchoulol production (Fig. 4E). A general, significant observation is the distinctly higher product yields in the dodecane layer of bisabolene compared to both sesquiterpene alcohols: 35.9 µmol * L^−1^ bisabolene equal 2.8 times and 6.1 times the molar titers of bisabolol (12.9 µmol * L^−1^) and patchoulol (5.9 µmol * L^−1^), respectively.

### Sesquiterpene production under small-scale High Density Cultivation (HDC) conditions

According to the data from both standard batch cultivation systems described above, cultures of high cell densities appear to be beneficial for high volumetric sesquiterpene product yields. Indeed, final bisabolene titers were even higher in the 6-well-plate experiment than in the MC system, which can be due to better light penetrance, higher mixing efficiency that favors cell-dodecane contacts, and higher evaporation-derived final cell densities. A recently developed HDC system allows for sustainably fast photoautotrophic growth, with a high potential for efficient biomass accumulation and high yields of natural products [28, 29]. In this study, we tested HDC for its value regarding heterologous terpenoid production. Strains P_*petE*__AgBIS, P_*petE*__*Mr*BBS, P_*petE*__PcPc and P_*petE*__mVenus were grown in nutrient-rich mineral CD medium including addition of 4 µM Cu^2+^ at tp 0h and 48h for stable induction of P_*petE*._. One of three independent cultivation experiments was extended to a total of eight days in a semi-batch mode: after 96h, the batch culture were 1:1 [v:v] replenished with fresh medium (Fig. 5A).

**Figure 5:**
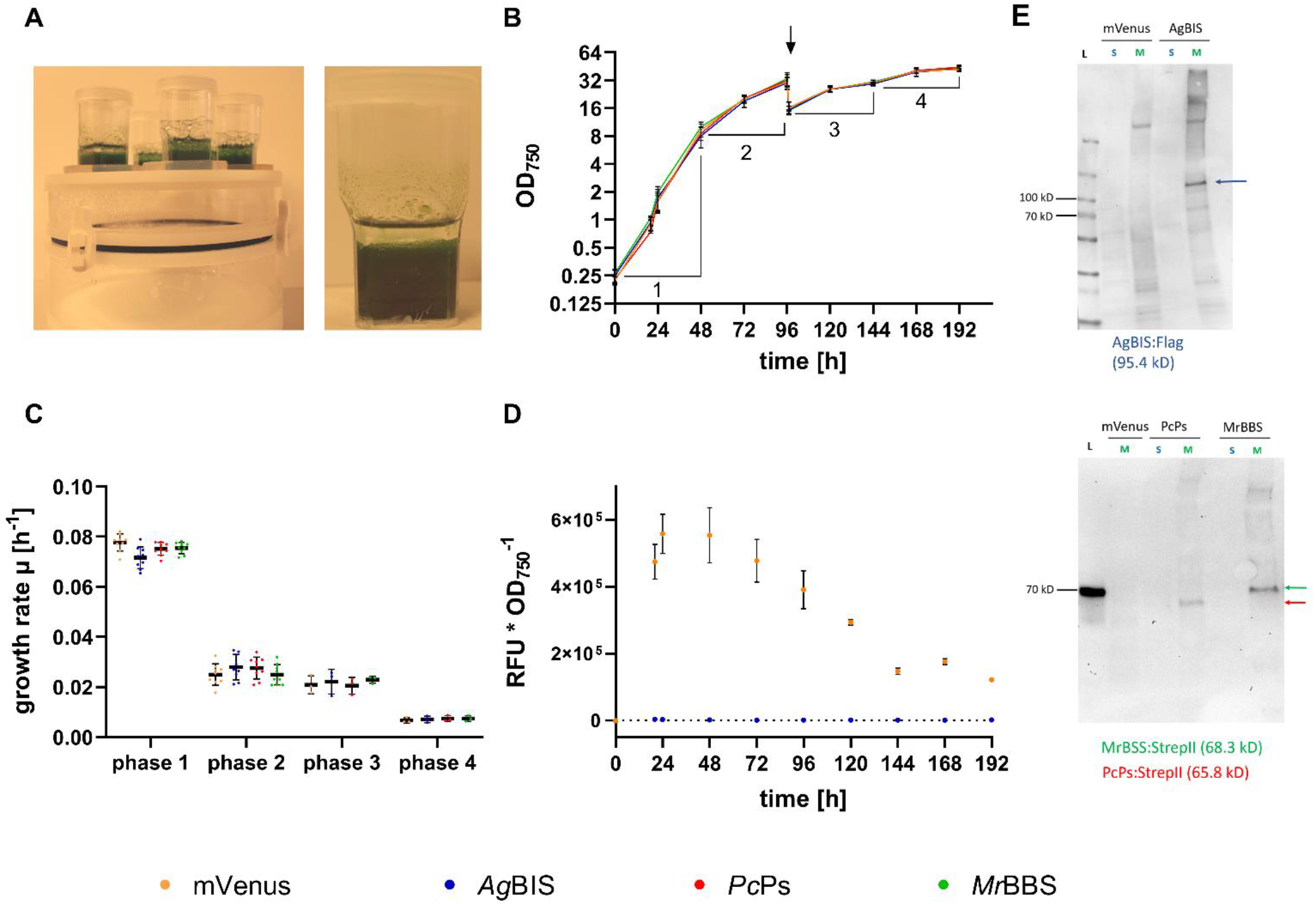
Growth characteristics and heterologous gene expression in HDC cultures. (A) HDC setup in HDC 6.10B system (Celldeg). Cultures were supplemented each with 25% [v/v] dodecane overlay and shaken at 320 rpm (4 mm orbit) with increasing light intensities. (B) In three independent cultivation runs, cell densities (OD_750_) of strains *P*_petE__*AgBIS*, *P*_petE__*MrBBS*, *P*_petE__*PcPc* and *P*_petE__mVenus were recorded daily over time periods of 2 x 4 days and 1 x 8 days. Data represent mean values from nine (tp 0, 24, 48, 96h) and three (tp 20, 97, 120, 144h, 192h) independent cultures with SD. The y-axis scale is logarithmic (log2). The arrow indicates the time point of media replenishment (96h). (C) Growth rate µ calculated from the OD_750_ data in B. Data represent single and mean values with SD. Phase 1 and 2 correspond to time periods 0-48h, 48-96h, 97-120h and 120-192h, respectively. (D) mVenus accumulation profile over time. Fluorescence data were divided by the corresponding OD_750_ data [λ Ex./Em. 485/535nm * OD_750_^−1^] and are depicted as mean values with SD (cf. B). (E) Immunoblot detection of *Ag*BIS:Flag (top), *Pc*Ps:StrepII and *Mr*BBS:StrepII (bottom) from HDC cultures harvested after 48h. Membrane fractions from P_*petE*__mVenus cultures were loaded as negative control. Apparent protein masses are estimated according to *PageRuler™ Prestained Protein Ladder* (L). The 70 kD marker is detected by the StrepII antibody (bottom). Theoretical masses of the proteins (calculated by the SnapGene software) are given in brackets. M, membrane fraction; S, soluble fraction; kD, kilo daltons.

The growth curves in Fig. 5B display bipartite growth profiles for all strains, with exponential phase 1 until 48h, and linear phase 2 until 96h, reaching final evaporation-corrected cell densities (OD_750_) of 30.67 ± 3.41 (P_*petE*__mVenus), 29.97 ± 4.16 (P_*petE*__*Ag*BIS), 32.03 ± 4.61 (P_*petE*__*Pc*Ps) and 33.26 ± 5.12 (P_*petE*__*Mr*BBS), respectively. The rather high standard deviations (SD) likely arise from varying evaporation rates during the cultivation runs (0.375 mL * d^−1^ vs. 0.5 mL * d^−1^), since the standard lids with inside oxygen filters were replaced with custom-made lids with outside filters. The variances for each replicate run were lower than for the total averages (table S4). One replicate cultivation was continued for 4 days after media replenishment, resulting in a bipartite growth pattern with linear phase 3 and stationary phase 4 (Fig. 5B). Final evaporation-corrected OD_750_ values after 192h were 41.92 ± 1.04 (P_*petE*__mVenus), 42.63 ± 2.53 (P_*petE*__*Ag*BIS), 44.34 ± 1.93 (P_*petE*__*Pc*Ps) and 43.67 ± 1.35 (P_*petE*__*Mr*BBS), respectively. Growth rates for the four growth phases are depicted in figure 5C and listed in table S4.

Specific accumulation of mVenus in P_*petE*__mVenus indicates growth-phase dependent variations in the activity of t P_*petE*_:BCD2-mediated gene expression under HDC conditions (Fig. 5D). The relative mVenus fluorescence peaks during exponential phase 1 at tp 24h and slightly decreases during linear phase 2 to ~70% at tp 96h, despite Cu^2+^ replenishment at tp 48h. After tp 96h in the extended semi-batch run, mVenus levels further decline to ~22% of the peak levels during phase 4. Here, downregulation of P_*petE*_ activity after transition into linear phase [49] and - potentially - a regulatory shift of the copper resistance system [50, 51] can be determining factors. However, the measured levels still represent considerable cellular amounts, since blue-light induced fluorescence in the cell suspensions is clearly visible to the naked eye (Fig. S2). We therefore assume sufficient expression of the heterologous genes over the whole cultivation period, taking into account that specific product formation rates might change (i.e. decline) as a function of P_*petE*_:BCD2-driven enzyme accumulation and lower protein stability (compared to mVenus).

The presence of the three terpene synthases *Ag*BIS:Flag, *Pc*Ps:Strep and *Mr*BBS:Strep was confirmed by immunoblot analysis of extract samples from tp 48h (Fig. 5E). Interestingly, all TS are exclusively located in the membrane fractions of the cell extracts, a phenomenon that was previously reported for eukaryotic triterpene cyclases [52, 53]. In line with the hydrophobic nature of terpenoids, this observation may inspire targeted enzyme engineering strategies as well as strategies towards the rational design of co-localized whole-pathway and pathway-transporter modules.

Intriguingly, the final volumetric titers of all three sesquiterpenes at tp 96h were more than one order of magnitude higher than at the same time point in the previous batch cultivation: 57.22 mg ± 10.18 * L^−1^, 24.51 ± 2.73 mg * L^−1^ and 9.86 ± 0.84 mg * L^−1^ for bisabolene, bisabolol and patchoulol, respectively (Fig. 6A-C). Already at tp 48h of HDC, the yields were at least twice as high as the final titers in the MC cultivation (tp 192h, cf. Fig. 4). In line with the previous observations, reduced growth rates (in phase 2) are accompanied with distinctly increasing volumetric production rates that appear to be largely stable for bisabolene and patchoulol from phase 2 to 4 (Fig. 6D, E), while even a significant increase in productivity was measured for bisabolol in phase 4 (Fig. 6F). It is of note that, compared to corresponding time points and growth phases in the MC cultivation (tp 96h and 144h), lower specific product accumulation of bisabolene (1.45 ± 0.66 mg * OD_750_^−1^ * L^−1^, Fig. 6G) and bisabolol (0.688 ± 0.34 mg * OD_750_^−1^ * L^−1^, Fig. 6I) was detected after phase 1 and 2. Specific patchoulol accumulation at tp 96h in HDC (0.17 ± 0.06 mg * OD_750_^−1^ * L^−1^, Fig. 6H) was comparable with specific titers at the same time point in the MC experiment, but distinctly lower when compared to the end of MC phase 2. While attempts to compare production dynamics under these different growth conditions might be delusive, the general observations highlight the adverse intracellular carbon partitioning characteristics towards biomass production during phases of high growth rates. After media replenishing of HDC cultures and their transition into phase 3, specific product titers clearly increased up to 2.2-fold (bisabolene, Fig. 6G), 1.5-fold (patchoulol, Fig. 6H) and 2.1-fold (bisabolol, Fig. 6I). While the culture dilutions themselves largely contribute to this effect, concomitantly stable volumetric productivities indicate a sustained carbon flux towards product formation when biomass accumulation rates slightly cease in phase 3. Sustained MEP pathway flux under stationary conditions was described for *E. coli* [54], and phototrophic patchoulol production in the green algae *C. reinhardtii* was shown to increase to some extent in the stationary phase [37]. As for cyanobacterial systems, the highest so far reported terpenoid yields using the example of isoprene were achieved in a 21-days cultivation experiment under a semi-batch regime with optimized carbon supply [13]. In that study, volumetric isoprene production rates increased in stationary phase with optical densities of >8 OD_750_ units. However, in our system, specific productivities clearly stall when culture growth heads from linear towards stationary characteristics between phases 3 and 4, while volumetric productivities stay quantitatively stable. In contrast to the isoprene study, rather linear growth in combination with high cell densities turn out to be most beneficial for our system. Similar correlations between growth phase and terpenoid accumulation have been observed in previous studies: heterologous production of both GPP-derived limonene (C10) and FPP-derived bisabolene in *Synechococcus* sp. PCC 7002 showed higher volumetric rates during exponential/linear growth over stationary phase [35]; for limonene production in *Synechocystis* similar observations were reported [18].

**Figure 6:**
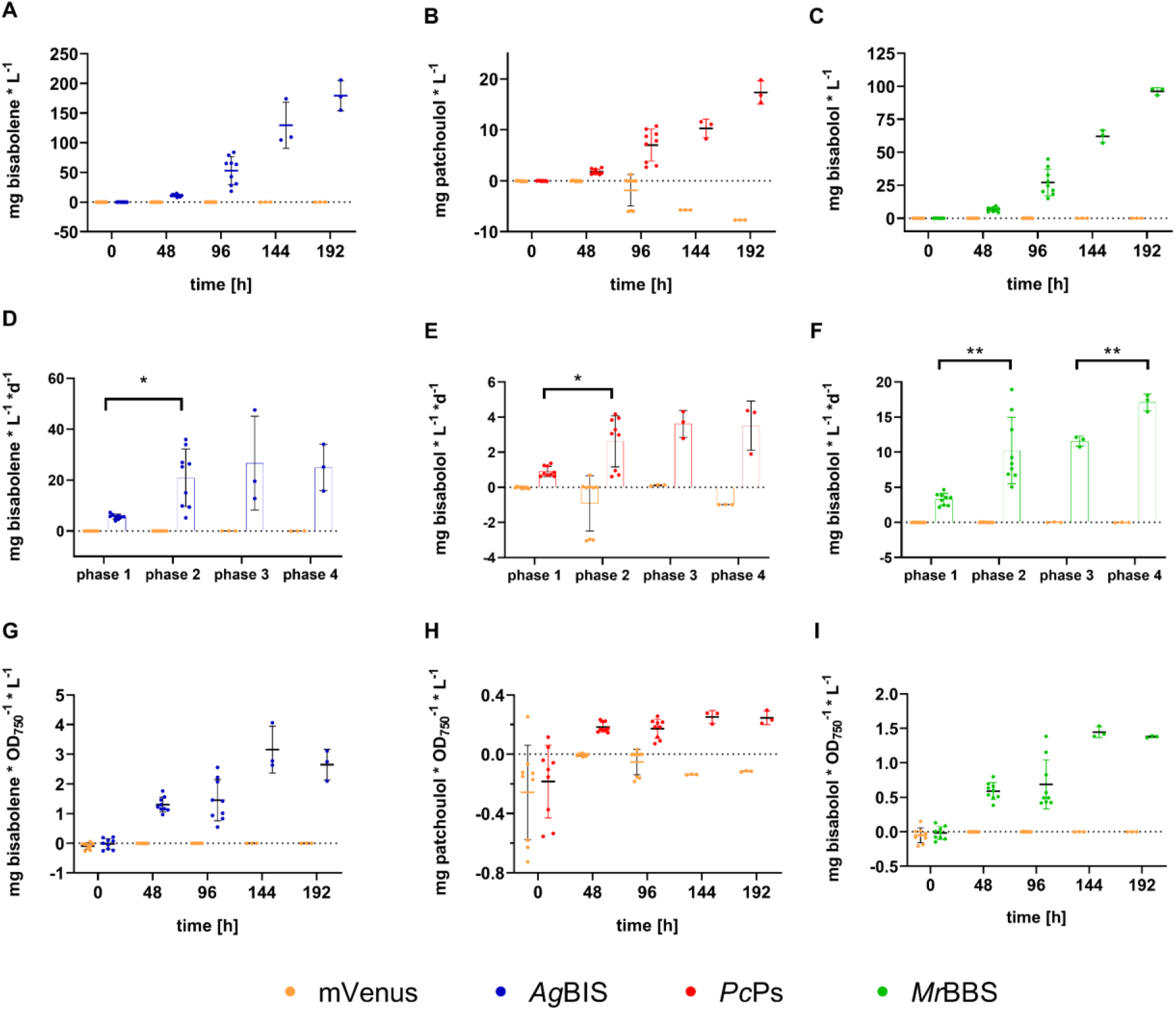
Time-dependent sesquiterpene accumulation profiles from HDC cultures. (A-C) Volumetric titers of bisabolene, patchoulol and bisabolol, respectively. Quantities were measured by GC-FID directly from the dodecane overlay and normalized to the total culture volume (mg * L^−1^). (D-F) Corresponding volumetric productivities calculated as the temporal yields (mg * L^−1^ * d^−1^) for growth phases 1 (0-48h), 2 (48-96h) 3 (97-120h) and 4 (120-192h), respectively. Statistical analysis was performed using the two-way ANOVA tool of the GraphPad Prism software (version 8.2.1.). *P < 0.05; **P< 0.01. (G-I) Specific product titers calculated as the quotient between volumetric titers and cell densities of the cultures (mg * L^−1^ * OD_750_^−1^).

Since fresh media supply (including fresh carbonate buffer for CO_2_ feeding) was introduced at the beginning of phase 3 in HDC, external nutrient availability is unlikely to play a key role for the characteristics of phases 3 and 4. Aside from light limitation [46], regulatory remodeling of nutrient acquisition at high densities is a conceivable scenario. During linear growth, a relative excess of carbon might be channeled into secondary metabolism, including the MEP pathway, while higher levels of self-shading would reduce the flux into the competing carotenoid biosynthesis branch. In microalgal systems the redirection of carbon flux from protein towards hydrocarbon biosynthesis under radical nutrient-depletion regimes was previously described and is exploited for triggering the (native) biosynthesis of cellular lipids [55, 56]. However, those metabolic routes do not involve the MEP pathway, and its activity in cyanobacteria as a function of nutrient availability needs to be elucidated. Aside from apparent physiological and metabolic shifts that occur in cultures of high cell densities, the total yields in the dodecane layer are likely improved according to an increase of the total interaction area between the entirety of the cell population and the organic phase. In a recent study on modular dynamics simulations of terpenoid extraction from membrane environments, Vermaas and co-workers [26] illustrated that hydroxylated terpenoids would cross the cellular membrane more slowly than more lipophilic compounds such as bisabolene.

This drawback is, however, energetically compensated by favorable characteristics of terpene alcohol adsorption by the dodecane-water interface. The authors further indicate that larger terpenoid compounds might rather get in direct contact with the dodecane layer, which is energetically advantageous for efficient extraction from the membrane bilayer. While general dodecane-mediated perturbance of the membrane structure might facilitate the desorption of smaller molecules as well, this aspect should be considered as part of the explanation why patchoulol with its more compressed tricyclic structure accumulated to ~4-6-fold lower levels than the structurally more extended bisabolol (Fig. 7C). Interestingly, the molar yield ratios between bisabolene and both sesquiterpene alcohols are higher during the productive phases at high densities (Fig. 7A, B), while a similar trend is also observable for the bisabolol:patchoulol ratio. This data indicate a higher benefit for non-oxygenated terpenoids from high cell densities in terms of extraction efficiencies. Likewise, the level of cyclization and molecular size seems to be relevant to extraction yields into the organic overlay, together suggesting that the selection of both the target compound and the corresponding extraction method are critical aspects to process design.

**Figure 7:**
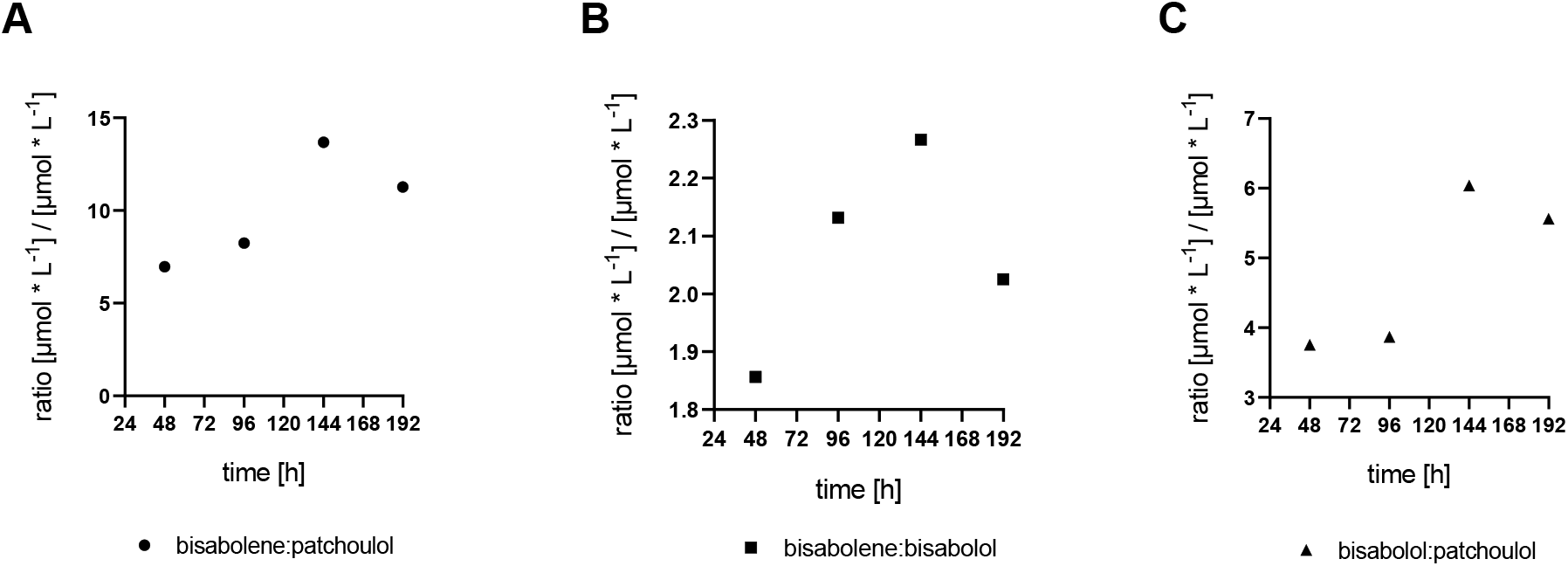
Molar yield ratios between the sesquiterpenoids over the HDC cultivation period. For calculation of yield ratios for the (A) bisabolene:patchoulol, (B) bisabolene:bisabolol and (C) bisabolol:patchoulol pairs, the mean volumetric titers [mg * L^−1^] were converted to molar titers [µmol * L^−1^] according to the molar mass of each compound: 204.357 g⋅mol^−1^ (bisabolene), 222.372 g⋅mol^−1^ (bisabolol) and 222.36 g⋅mol^−^ (patchoulol), respectively. Ratios for each time point were calculated by division.

Aside from the impact of diverging extraction efficiencies, general yield bias can result from intrinsic differences like enzyme activities and product toxicity. Bisabolol product toxicity was reported for *E. coli* [38], and might play a role in cyanobacterial systems too. However, no significantly consistent effects of any of the tested terpenoid products on cell viability and growth was observed in this study. While direct removal of the terpenoid by the dodecane sink would reduce potential intracellular toxicity-related effects in our system, a systematic dose-response study including external treatment will be required to gain a quantitative estimation of this aspect.

In summary, HDC does not improve specific productivity of the cell cultures for any of the sesquiterpenoids during high growth rates; specific titers rather stall in all strains during phases 1 and 2 (Fig. 6 D-F). Evidently, under constantly high growth rates biomass production vastly outcompetes terpenoid biosynthesis, clearly pointing at carbon partitioning accounting for the major bottleneck in efficient production. On the other hand, volumetric and specific titers of both bisabolene and bisabolol in semi-batch mode outcompete those from MC cultivation. Therefore, HDC provides an efficient platform for transferring cyanobacterial cultures into a beneficial state for high total product yields within short times periods. Particularly, the data after media replenishment suggest technological benefits from coupling HDC to a semi-continuous or continuous cultivation mode, which could facilitate sustainably high temporal yields over extended cultivation periods. Long-term studies using HDC reactors of higher volumetric scale will be important to further assess the potential of this technology for industrial-scale terpenoid production. Furthermore, dynamic metabolome studies of different growth phases under HDC conditions are highly desirable to shed further light on the reprogramming of carbon fluxes during these physiological transitions.

## Conclusions

We demonstrate the applicability of a commercial High Density Cultivation platform for efficient production of sesquiterpenoids in *Synechocystis*. The small-scale system can be used for *in situ* extraction protocols that involve organic overlays like dodecane and facilitates high-yield production within short time periods, making HDC certainly attractive for rapid screening procedures. Growth-phase related analysis further suggests a combination of linear growth and high cell densities as favorable conditions for cyanobacterial terpenoid biosynthesis with *in situ* harvest. The extraction gains are particularly pronounced for the non-oxygenated hydrophobic molecule bisabolene; but also yields of both oxygenated products are distinctly improved. In line with the different extraction yields, diverging physicochemical and structural properties of candidate terpenoids should be taken into account, when selecting a chemical of interest. This study further provides a promising outlook on emerging bioreactor designs – like e.g. solid-phase biofilm reactors - that inherently resort to high-density strategies.

## Material and Methods

### Plasmid construction and transformation

The common vector was the conjugative plasmid RSF1010-based pSHDY*in, which is a derivative of pSHDY (version w/o *mobA*Y25F point mutation) [41] with optimized insulation of the default expression cassette. The latter consists of the copper-inducible promoter P_petE_ (from *Synechocystis*), the 5’UTR BCD2 [44], a codon optimized coding sequence (CDS) for the mVenus fluorophore [40] including a second stop codon and the rrnB T1/ T7Te double terminator (<u>BBa_B0015</u>). To insulate target gene expression from adjacent antibiotic cassettes, a 200-bp “non-sense” sequence according to Yeung et al. [57] preceded by the rrnB T1/ T7Te double terminator was inserted between prefix and (upstream of the) promoter. The vector backbone harbors streptomycin/ spectinomycin (Sm^R^) and kanamycin (Kan^R^) resistance cassettes upstream and in divergent orientation of the insulated promoter region (Fig. S1A).

For expression of *Ag*BIS, *Mr*BBS and *Pc*Ps, the corresponding CDS were codon optimized for expression in *Synechocystis* using DNA2.0’s GeneDesigner (*Ag*BIS) and IDT’s codon optimization tool (*Mr*BBS, *Pc*Ps, mVenus), respectively. For modular cloning 2^*nd*^ and 3^rd^ codons were transformed into an NheI restriction site (*Mr*BBS, *Pc*Ps, mVenus) or an (NheI-compatible) SpeI site (*Ag*BIS). *Ag*BIS was C-terminally fused to a 1x Flag tag via an RSGSGS linker (RGS); *Mr*BBS and *Pc*Ps were fused to a Strep tag II via a GSGSGS linker (GS) on the C-terminus. A second Stop codon was added to the CDS of mVenus, *Mr*BBS-GS-StrepII and *Pc*Ps-GS-StrepII. Downstream of the full CDS and rrnB T1/ T7Te double terminators a PstI site was inserted for common NheI(SpeI)/PstI-mediated replacement of the expression units (Fig.S1B-D).

Primers and synthetic dsDNA parts are listed in table S1. All plasmids are listed in table 1 and will soon be accessible at the Addgene Repository under the following accession numbers: 133970 (pSHDY*in_PpetE:BCD2:mVenus), 133971 (pSHDY*in_PpetE:BCD2:AgBIS:Flag), 133972 (pSHDY*in_PpetE:BCD2:PcPs:Strep) and 133973 (pSHDY*in_PpetE:BCD2:MrBBS:Strep). Full plasmid sequences are further available on Figshare: 10.6084/m9.figshare.10265072 (mVenus), 10.6084/m9.figshare.10265063 (*Ag*BIS), 10.6084/m9.figshare.10265102 (*Mr*BBS), 10.6084/m9.figshare.10265069 (*Pc*Ps)

**Table 1:**
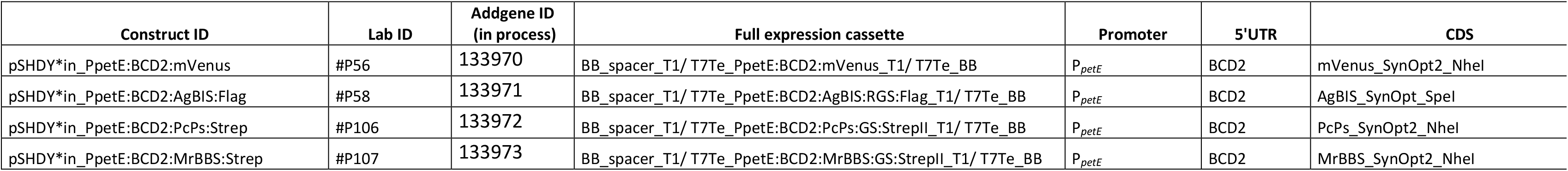
Genetic constructs for heterologous expression of mVenus and sesquiterpenoid synthases in *Synechocystis*.

### Strains and cultivation conditions

*Escherichia coli* strains DH5α [58] and HB101 (harboring conjugal plasmid pRL443) [59] were used for cloning and triparental mating, respectively. All *E. coli* strains were cultivated on plant-based LB-agar plates or in liquid LB medium (lysogeny broth) at 37 °C. Antibiotics were added to *E. coli* culture medium to the final concentrations of 20 μg mL^−1^ spectinomycin (spec), 100 μg mL^−1^ ampicillin (amp) and 50 μg mL^−1^ kanamycin (kan), respectively.

*Synechocystis* sp. PCC 6803 strain GT-U, a derivative of the non-motile, glucose-tolerant strain GT-Kazusa [60] was used as final host for all conjugations. For selection and strain maintenance *Synechocystis* was cultivated on 0.75% [w/v] agar (Type A, Sigma) plates containing TES-buffered (10 mM, pH 8.0) BG-11 mineral medium [61]. Pre-cultures were grown in TES-buffered BG-11 liquid medium in 6-well polystyrene at 30 °C and 120 rpm. By default, CuSO_4_ and Co(NO_3_)_2_ was not added to any basic BG11 media preparations in this study in order to: (i) minimize uncontrolled Cu^2+^ induction of P*petE*, and to (ii) provide reference data for follow-up studies using Co^2+^-induction. For contamination control and plasmid maintenance, all cultures of transformed *Synechocystis* strains were supplied with 20 μg mL^−1^ spectinomycin and 50 μg mL^−1^ kanamycin.

- **Experimental small-scale cultivation in 6 well plates** Screening assays of sesquiterpene producer strains were performed in 6 well polystyrene cell culture plates, with total culture volumes of 3 mL BG11 including both antibiotics. The cell densities of each six preculture batches per strain were adjusted to OD_750_ 1.0 by appropriate media-dilution with fresh BG11. Each adjusted cell suspension was distributed over two wells, resulting in a total of twelve batches per strain. Each six batches were supplemented with 2 µM CuSO_4_ for P_*petE*_induction (+Cu), while each six control batches were grown in copper-free medium (-Cu). An overlay of 600 µL dodecane was added to all cultures. Cultivation took place on a ‘Standard analog shaker, Model 5000 (VWR; orbit: 25 mm; frequency: 120 rpm) at 30 °C and under constant illumination with fluorescent white light at ~100 µmol photons * m^−2^ * s^−1^. OD_750_ and mVenus fluorescence were monitored at tp 0h, 24h, 48h and 96h (table S2). After 4 days, each 198 µL of dodecane were sampled for GC-FID and GC-MS analysis. Due to high evaporation losses of culture media, with a calculated evaporation rate of 0.1 mL * d^−1^, all calculations were corrected for this factor. All data from this experiment are compiled in table S2.
- **Medium-scale cultivation in Multi-Cultivator (MC)s** For cultivation in the Multi-Cultivator system MC 1000 (Photon System Instruments), pre-cultures were centrifuged for 5 min at 2500 g and 22 °C and resuspended in bicarbonate-enriched BG11 (50 mM NaHCO_3_; 50 mM TES) including spectinomycin and kanamycin with a total volume of 60 mL. Thereby, the initial cell densities were adjusted to OD_750_ ~0.25. Each strain was cultivated in quadruplicate from the same pre-culture. The cultivation vessels were distributed over two separate MC devices, at which duplicates of each strain were separated by each three slots per device to minimize position-related effects. Strains were illuminated with the “cool white light” LED version at initial intensities of 50 µmol photons * m^−2^ * s^−1^, which was increased to 70 µmol photons * m^−2^ * s^−1^ after six days. Temperature control was set at 30 °C, resulting in small actual temperature fluctuations between 29 °C and 32 °C during the cultivation period. All cultures were supplied with a manually adjusted, moderate and constant air flow provided by the supplier’s aeration pump. An overlay of 3 mL dodecane was added to all cultures. For on-site culture sampling, silicone tubes were inserted into the vessels through the outflow tubing, locked by plastic clamps and connected to 3 mL syringes through female Luer adapters. For daily monitoring of OD_750_ and mVenus fluorescence, each ~0.5 mL dead volume and ~0.5 mL sample volume was removed from the cultures. At time point 6d, additional 5 mL culture for Immunoblot analysis were removed before replenishing all cultures with 12 mL of fresh bicarbonate-enriched BG11. For GC-FID measurements of terpenoids, at time points 0h, 48h, 96h, 144h and 192h each 198 µL dodecane were removed from the overlay, which was replenished with the same amount of fresh dodecane. All data from this experiment are compiled in table S3.
- **High-density cultivation (HDC)** HDC was performed in three separate runs using the HDC 6.10 starter kit (Celldeg), which uses carbonate buffer as CO_2_ source for 10 mL cultivator vessels. The buffer reservoir was filled with 200 mL of a 3M KHCO_3_/3M K_2_CO_3_ (volume ratio 9:1) solution to provide a CO_2_ partial pressure of 90 mbar (reference T = 20 C°), according to the manufacturer’s recommendation. The exact nutrient-enriched media composition (CD medium) is accessible on protocols.io (dx.doi.org/10.17504/protocols.io.2bxgapn). Like BG11, basic CD media preparations did not contain CuSO_4_ and Co(NO_3_)_2_ in this study. For inoculation, pre-cultures were grown in BG11 (-Cu) ~ and centrifuged for 5 min at 2500 g and 22 °C before cell pellets were resuspended in 8 mL CD medium including spectinomycin and kanamycin. Thereby, the initial cell densities were adjusted to OD_750_ ~0.25. Each culture was supplied with 4 µM CuSO_4_ at tp 0h and 48h. An overlay of 2 mL dodecane was added to all cultures. In one replicate run (run 3), all cultures were replenished with 1 Vol. fresh CD medium (4 mL culture + 4 mL medium) at tp 96h, including 2 µM CuSO_4_ (final concentration). These cultures were further treated with 4 µM CuSO_4_ at tp 144h. Cultivation took place at 30 °C under constant, multi-directional illumination with fluorescent white light in ‘Versatile Environmental Test Chamber (Sanyo) w/o humidifier. The sequence of increasing light intensities was: 250 µmol photons * m^−2^ * s^−1^ (tp 0h-24h), 490 µmol photons * m^−2^ * s^−1^ (tp 24h-48h), 750 µmol photons * m^−2^ * s^−1^ (tp 48h-192h). Cultures were constantly shaken at 320 rpm on an IKA KS 130 basic orbital shaker (ø = 4 mm). For measurements of OD_750_ and mVenus fluorescence, culture volumes between 5 µl and 100 µL were sampled in dependence on the respective cell density. At tp 48h of run 1, additional 200 µL were removed for Western Blot analysis and replenished with fresh medium. For GC-FID measurements of terpenoids, at time points 0h, 48h and 96h each 198 µL dodecane were removed from the overlay and replaced with the same amount of fresh dodecane. Cultures of run 3 were further samples at tp 144h and 192h. All data from this experiment are compiled in table S4. A detailed protocol is accessible on protocols.io (dx.doi.org/10.17504/protocols.io.757hq9n).

### *In-vivo* fluorescence and OD_750_ measurements

Measurements of optical cell density (OD) and *in vivo* mVenus fluorescence in *Synechocystis* was conducted in ‘TC Plate 96 Well, Standard, F’ (Sarstedt) using the “Plate Chameleon V Microplate Reader” (Hidex). OD was defined as the absorbance at 750 nm, fluorescence was measured using the 485 nm excitation filter and 535 nm emission filter. All data was corrected for the optical length path (factor 5) and the dilution factor of each sample (standard cultivation: factor 5; HDC: factor 2-40, depending on cell density). All fluorescence and OD_750_ values were adjusted by subtraction of the corresponding blank media values. For calculating relative fluorescence levels, adjusted fluorescence values were divided by the adjusted OD_750_ values.

### Immunoblot analysis

For extract preparation each 4 mL of cell cultures from tp 48h of HDC run 1 were spun down for 10 min at 4000 g and 4 °C (Centrifuge 5804R, Rotor A-4-44, Eppendorf). Pellets were resuspended in fresh 1 mL ‘thylakoid buffer’ (50 mM HEPES-NaOH, pH 8.0; 5 mM MgCl_2_, 25 mM CaCl_2_; 50 mM Na_2_-EDTA, pH 8.0; 1 x ProteaseArrest™ agent, G-Bioscience). Equivalents of each 50 OD units were 5x diluted in thylakoid buffer to a final volume of 1 mL. A volume of ~200 µL 425-600 µm glass beads (acid-washed, 20-40 U.S. sieve, Sigma) was added and samples were homogenized using a Precellys24 tissue homogenizer (Bertin technologies) for 2 x 30 s at 5600 rpm with a 2 min interval. For separation into soluble and membrane fractions, the crude extracts were transferred into a fresh tube and centrifuged for 30 min at 15000g and 4 °C (Centrifuge 5427R, Eppendorf). The soluble supernatant (blue color) was transferred to a fresh tube; the insoluble pellet (green color) was resuspended in 50 µL thylakoid buffer including 0.1% [v/v] Triton-X 100. For protein quantification, duplicates of each 2.5 µL sample were analyzed with the DC™ protein assay kit (Biorad) in 96-well plates, following the manufacturer’s instructions. A linear standard curve of BSA (0, 0.2, 0.4, 0.8, 1.6 mg * mL^−1^) was measured in parallel. The colorimetric readout was determined at 750 nm in a “Plate Chameleon V Microplate Reader” (Hidex).

For SDS PAGE, samples of each 20 µg protein were mixed with ¼ volume of 4x loading dye (400 mM DTT; 250 mM Tris-HCl, pH 6.8; 40% [w/v] glycerol; 8% [w/v] SDS) and denatured for 30 min at 50 °C. Samples were loaded onto Mini-Protean TGX stain-free 4-15% gradient gels (Biorad) and separated under a constant voltage of 200 V. PageRuler™ Prestained Protein ladder (Thermo Fisher) was used as molecular weight reference. For rapid transfer of separated proteins to a PVDF membrane, the ‘Trans-Blot Turbo Transfer System’ (Biorad) was used in combination with the ‘Trans-Blot Turbo Transfer Pack’ (Biorad). The transfer proceeded for 30 min at 25 V. Membranes were rinsed in TBS-T buffer (100 mM Tris-HCl, pH 7.5; 150 mM NaCl; 0.1% [v/v] Tween 20) and incubated o/n at 4 °C in soy milk (Alpro), before they were washed three-times for 15 min in TBS-T at 90 rpm and RT.

For Flag-tag detection membranes were incubated for two hours with the monoclonal anti-Flag M2 antibody from mouse (F3165, Sigma) in a 1:5000 dilution (in TBS-T) at RT under constant agitation. A 1:5000 dilution of the “Immun-Star Goat Anti-mouse-HRP conjugate” (1705047, Biorad) was used as secondary antibody under the same conditions. The same protocol was applied for Strep-tag detection using a 1:1000 dilution of the anti-Strep-tag II antibody (ab76949, abcam) and a 1:10000 dilution of ‘Immun-Star Goat Anti-rabbit-HRP conjugate’ (170-5046, Biorad) as secondary antibody. After each antibody treatment, membranes were washed three times for 15 min in TBS-T at 90 rpm and RT. Chemiluminescent detection of the HRP-conjugates was performed with the ‘Clarity Western ECL’ kit (Biorad), using ChemiDoc XRS system (Biorad) with cumulative image recording using the Quantity One software (Biorad).

### GC-FID detection of sesquiterpenes and sesquiterpene alcohols

The 198-µL dodecane overlay fractions were collected in 1.5 mL clear glass GC vials with a 9 mm Silicone/PTFE closure (VWR) and supplied with each 2 µL ß-caryophyllene (BCP, Stock: 25 µg * µL^−1^) as internal standard (IS). Each1 µL sample was injected from an autosampler into the Perkin Elmer GC 580 system, equipped with Elite-Wax Polyethylene Glycol Series Capillary (Perkin Elmer, length: 30m; inner diameter: 0.25 mm; film thickness: 0.25 µM) and flame ionization detector (FID). The system operated with N_2_ as carrier gas with a flow rate of 50 mL *min^−1^ and the following temperature profile: injection temperature = 250 °C, 1 min 100 °C, ramp 5 °C * min^−1^ to 160 ͦC, 2 min hold, ramp 10 °C min^−1^ to 240 °C, flow rate = 50 mL *min^−1^.

A detailed protocol is accessible on protocols.io (dx.doi.org/10.17504/protocols.io.kj2cuqe)

### Quantification of sesquiterpenes and sesquiterpene alcohols

For quantitative analysis calibration curves were obtained with commercial references of (*E*)-α-bisabolene (A18724, Alfa Aesar), (-)-α-bisabolol (95426, Sigma) and patchouli alcohol (5986-55-0, Sigma) as external standards (ES). β-caryophyllene (BCP, ≥80%, W225207, Sigma) was used as an internal standard (IS) in a total concentration of 2.5 µg * mL^−1^. Terpenoid yields were calculated based on GC-FID data as product titer in the culture [mg L^−1^], as cell-density normalized specific titer (mg L^−1^ OD^−1^):

For peak area [µV⋅s] normalization individual values were divided by the corresponding IS peak area and multiplied with the mean IS value of all biological and ES samples. Raw volumetric concentrations [µg * mL^−1^] were calculated with the linear equation from ES calibration charts derived from the same GC cycle. For bisabolene, each value was further multiplied with factor 0.2752, to correct for impurities of the commercial standard [36]. The mean raw concentration of all blank samples from the same run were subtracted from each value. The total concentration in the dodecane overlay was extrapolated with the corresponding scale factor and corrected for the addition of each 2 µL IS to 198 µL sample (e.g. (10 µg * mL^−1^ * 2/1000)*(200/198) = 0.02 mg * 2 mL^−1^ overlay). To correct for dodecane removal at previous time points, the concentration (0.02 mg * 2 mL^−1^) was multiplied with the total overlay volume (e.g. 2 mL) and divided by the total volume minus the total of removed volume (e.g. (0.02 *2)/(2-0.198) = 0,022 [mg * 2 mL^−1^]. This value was divided by the actual culture volume and multiplied with factor 1000 to extrapolate the final product titer [mg * L^−1^].

### GC-MS detection and of sesquiterpenes

Gas chromatography mass spectroscopy (GC-MS) analysis of dodecane overlay samples was conducted as previously described [36, 37]. Attribution of detected peaks to sesquiterpenoid compounds was performed by comparison of retention times and mass fractionation patterns (m/z spectra) with authentic standards.

## Author contributions

DD and PL conceptualized the study and conducted experiments. DD analyzed and interpreted data and drafted the manuscript. JW conducted mass spectrometry analysis and data interpretation. OM established the HDC platform and performed experiments. JR implemented the analytical pipeline and conducted experiments. All authors reviewed and approved the manuscript.

## Acknowledgements

This work was supported by ÅForsk foundation (grant no. 16-538), and by the NordForsk NCoE program “NordAqua” (project number 82845). We thankfully acknowledge Lars Bähr and Arne Wüstenberg (Celldeg GmbH, Berlin) for productive technical discussions and custom-made product design. Thanks to Sara Nilsson (Uppsala University) for establishing the bisabolene analytics protocol at Ångström Laboratory. Thanks to Ilka M. Axmann for kindly providing the pSHDY backbone. Further thanks to Kyle J. Lauersen (KAUST, Thuwal) and Paul Hudson (KTH, Stockholm) for valuable discussions and inspirations. This work has further been supported by the technology platform and infrastructure at the Center for Biotechnology (CeBiTec) at Bielefeld University (JW).

