## Supplemental Figures S1 and S2 for "High density cultivation for efficient sesquiterpenoid biosynthesis in *Synechocystis* sp. PCC 6803"

### Supplementary Figures

### A

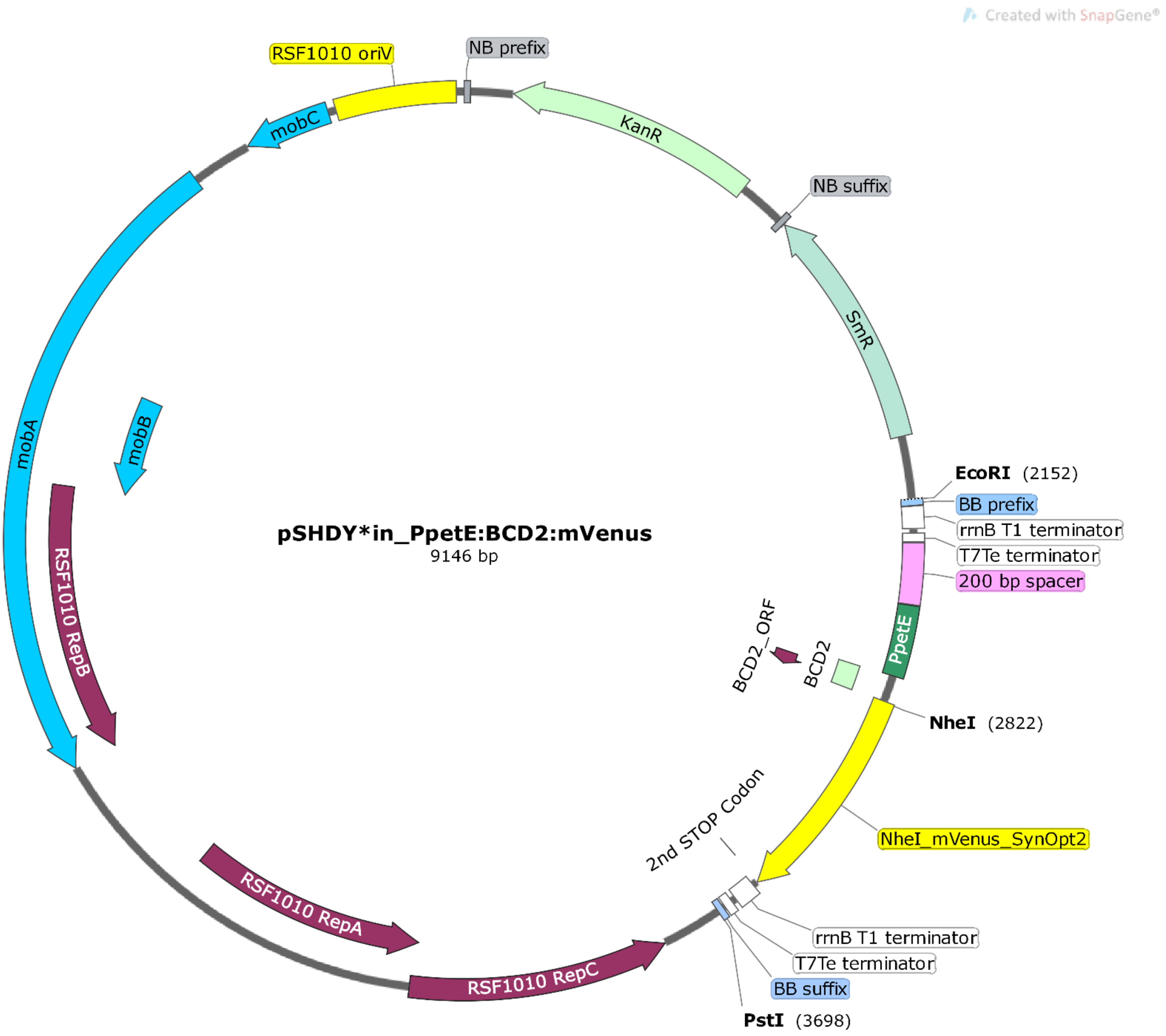

**B**

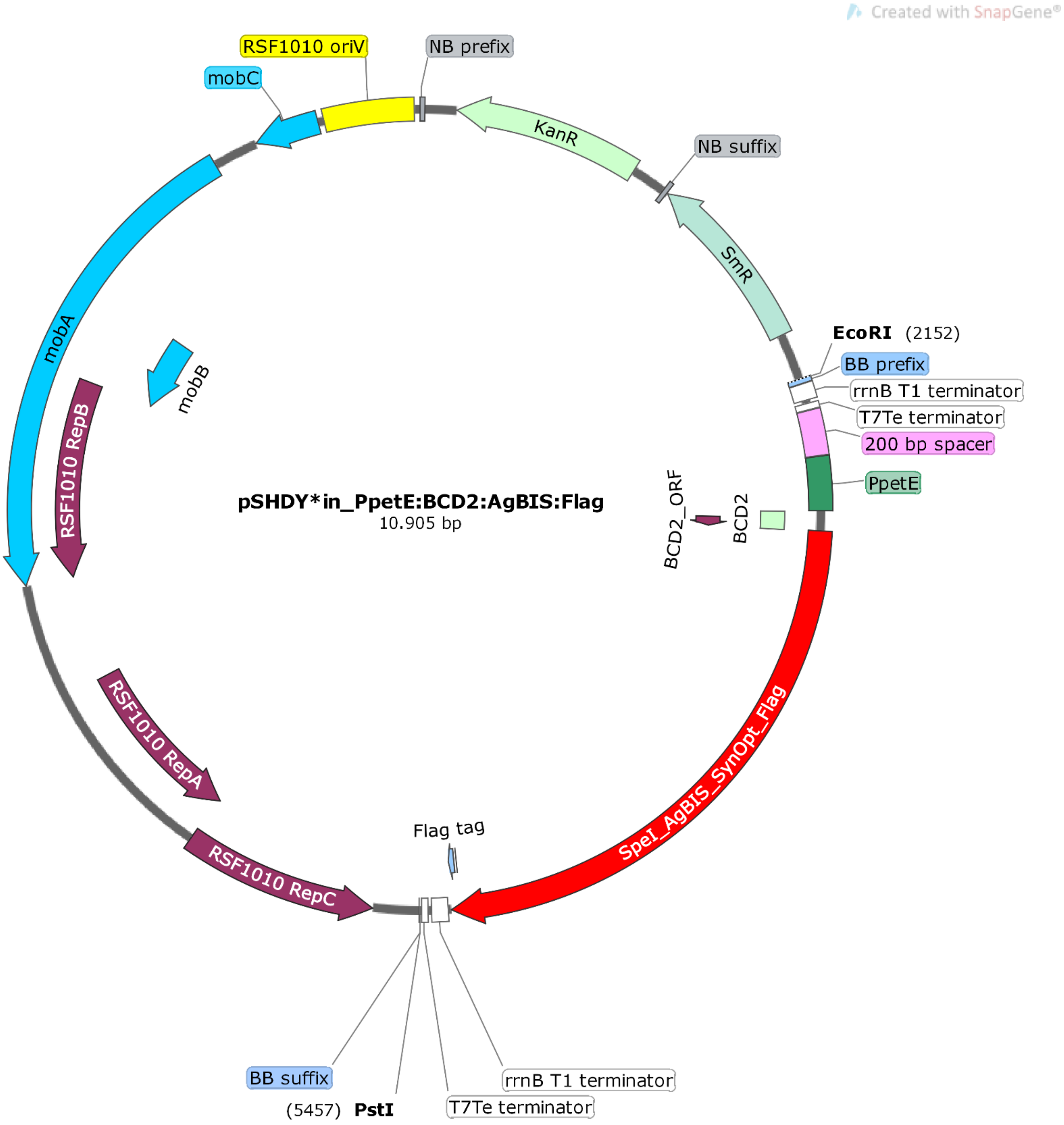
Created with SnapGene®
RSF1010 oriV
NB prefix
mobC
KanR
NB suffix
SmR
EcoRI (2152)
BB prefix
rrnB T1 terminator
T7Te terminator
200 bp spacer
PpetE
mobA
mobB
RSF1010 RepB
pSHDY*in_PpetE:BCD2:AgBIS:Flag
10.905 bp
BCD2_ORF
BCD2
SpeI_AgBIS_SynOpt_Flag
RSF1010 RepA
Flag tag
RSF1010 RepC
BB suffix
(5457) PstI
rrnB T1 terminator
T7Te terminator

C

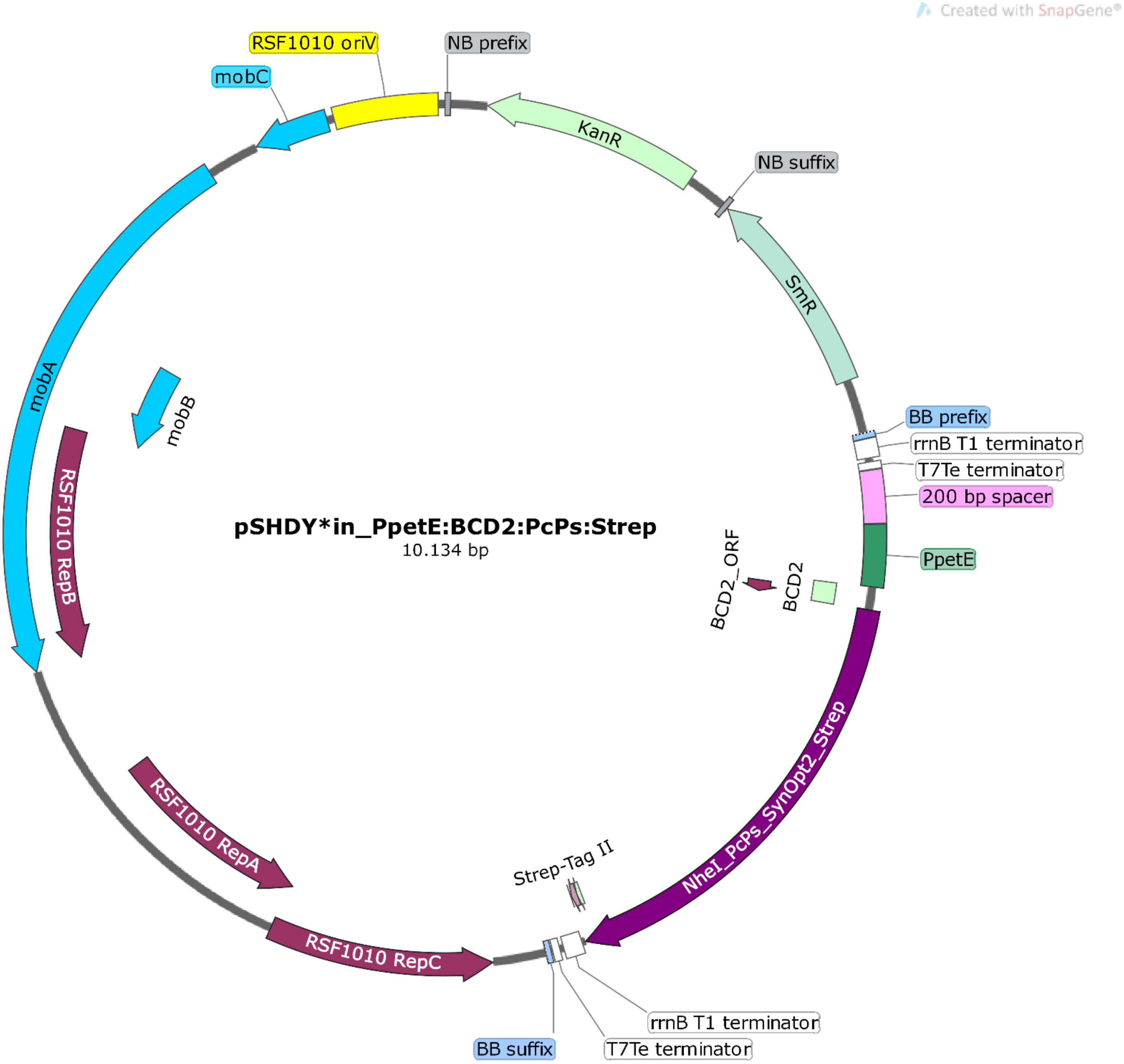
Created with SnapGene®
pSHDY*in_PpetE:BCD2:PcPs:Strep
10.134 bp
RSF1010 oriV
NB prefix
mobC
KanR
NB suffix
SmR
BB prefix
rrnB T1 terminator
T7Te terminator
200 bp spacer
PpetE
BCD2_ORF
BCD2
NheI_PcPs_SynOpt2_Strep
Strep-Tag II
rrnB T1 terminator
T7Te terminator
BB suffix
RSF1010 RepC
RSF1010 RepA
RSF1010 RepB
mobA
mobB

**D**

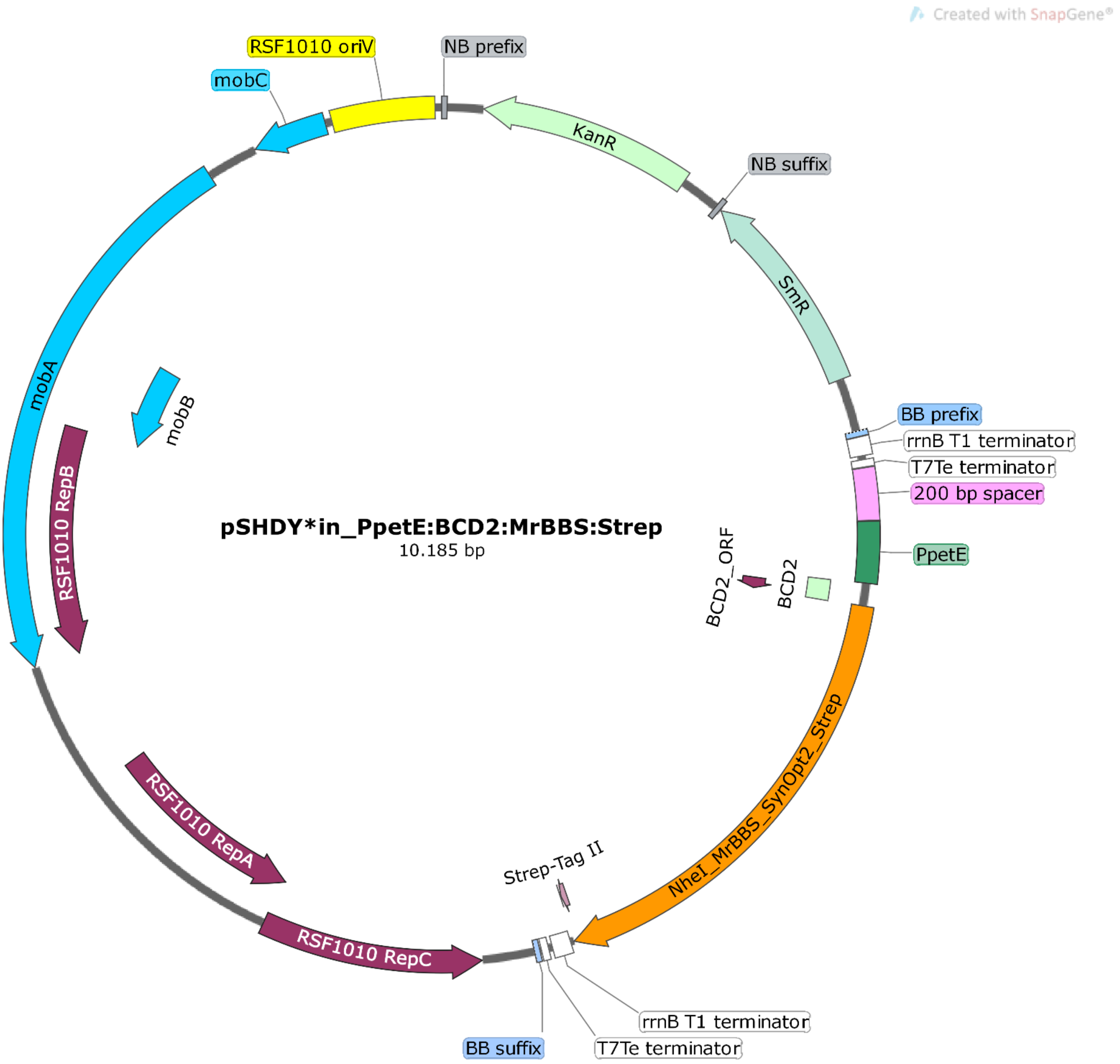

Created with SnapGene®
RSF1010 oriV
NB prefix
mobC
KanR
NB suffix
SmR
mobA
mobB
BB prefix
rrnB T1 terminator
T7Te terminator
200 bp spacer
pSHDY*in_PpetE:BCD2:MrBBS:Strep
10.185 bp
RSF1010 RepB
BCD2_ORF
BCD2
PpetE
NheI_MrBBS_SynOpt2_Strep
RSF1010 RepA
Strep-Tag II
RSF1010 RepC
rrnB T1 terminator
BB suffix
T7Te terminator

***Figure S1:*** Map of vectors (A) pSHDY*in_PpetE:BCD2:mVenus, (B) pSHDY*in_PpetE:BCD2:AgBIS:Flag, (C) pSHDY*in_PpetE:BCD2:PcPs:Strep and (D) pSHDY*in_PpetE:BCD2:MrBBS:Strep used for reporter fluorescence monitoring (A) and heterologous biosynthesis of sesquiterpenes (B, C, D) in *Synechocystis*, respectively.Insertion of the constructs ‚MrBBS:GS:Strep: T1/ T7Te ‘ and ‚PcPS:GS:Strep: T1/ T7Te ‘ into the pSHDY_in_PpetE:BCD2 backbone was conducted via NheI ($2^{nd}/3^{rd}$ codon of CDS) and PstI (downstream of double terminator rrnB T1/ T7Te), using pSHDY-in_PpetE:BCD2:mVenus as cloning template. Construct ‚AgBIS:AGS:Flag:T1/ T7Te‘ was inserted via SpeI and PstI, thereby destroying the compatible NheI site in the template. Gene expression in *Synechocystis* was mediated by the native copper-inducible promoter P*petE* and the 5’UTR BCD2, harboring an insulated translation initiation feature [1]. Transcription terminates at the double terminator B0015 (rrnB T1/ T7Te). For improved insulation of the $P_{petE}$:BCD2 expression module, B0015 and a *non-sense* 200-bp spacer [2] were ligated upstream of $P_{petE}$. KanR, kanamycin resistance cassette; SmR, streptomycin/spectinomycin resistance cassette; Rep, genes encoding replication proteins; *mobABC*: genes encoding mobilization proteins required for triparental mating; BB, biobrick; NB, neobrick.

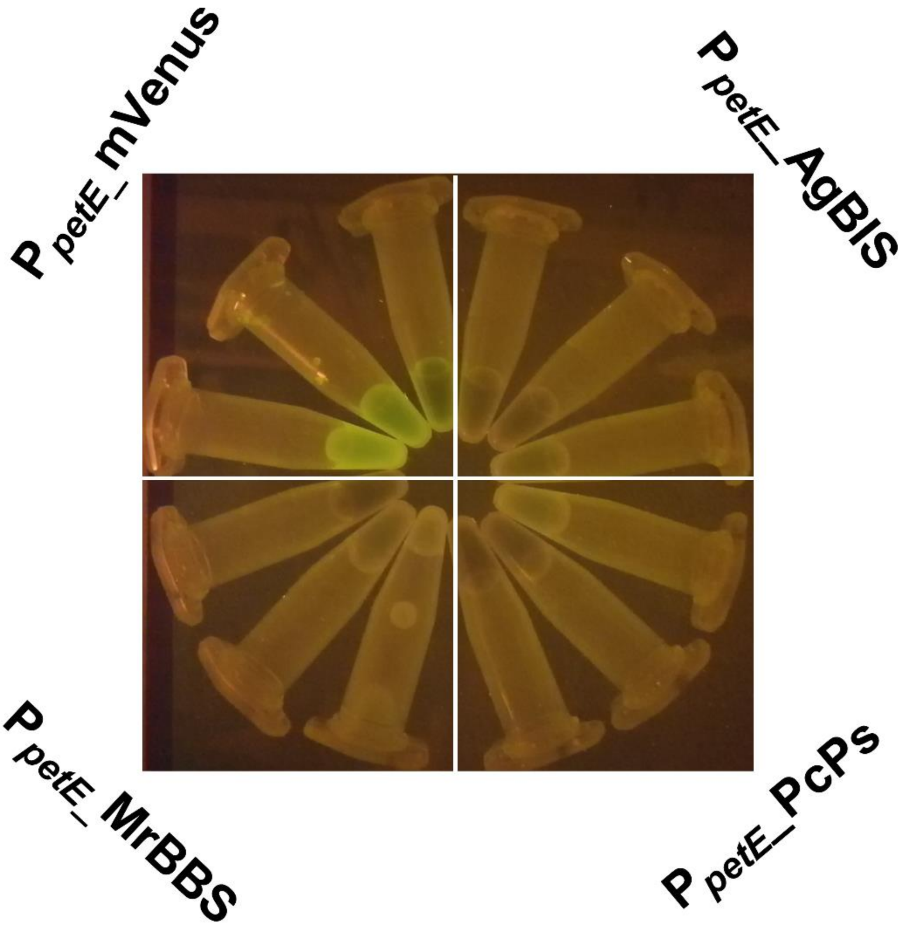


**Figure S2:** Fluorescence of *Synechocystis* strains at tp 192h of HDC. Aliquots of the cell suspensions were subjected to blue light (470 nm) illumination on a “Safeview-Mini2” transilluminator.

1. Mutalik VK, Guimaraes JC, Cambray G, Lam C, Christoffersen MJ, Mai QA, Tran AB, Paull M, Keasling JD, Arkin AP *et al*: **Precise and reliable gene expression via standard transcription and translation initiation elements**. *Nat Methods* 2013, **10**(4):354-360.
2. Yeung E, Dy AJ, Martin KB, Ng AH, Del Vecchio D, Beck JL, Collins JJ, Murray RM: **Biophysical Constraints Arising from Compositional Context in Synthetic Gene Networks**. *Cell Syst* 2017, **5**(1):11-24 e12.
